# Bioorthogonal protein-DNA conjugation methods for force spectroscopy

**DOI:** 10.1101/631572

**Authors:** Marie Synakewicz, Daniela Bauer, Matthias Rief, Laura S. Itzhaki

## Abstract

Accurate and stable site-specific attachment of DNA molecules to proteins is a requirement for many single-molecule force spectroscopy techniques. The most commonly used method still relies on maleimide chemistry involving cysteine residues in the protein of interest. Studies have consequently often focused on model proteins that either have no cysteines or with a small number of cysteines that can be deleted so that cysteines can then be introduced at specific sites. However, many proteins, especially in eukaryotes, contain too many cysteine residues to be amenable to this strategy, and therefore there is tremendous need for new and broadly applicable approaches to site-specific conjugation. Here we present bioorthogonal approaches for making DNA-protein conjugates required in force spectroscopy experiments. Unnatural amino acids are introduced site-specifically and conjugated to DNA oligos bearing the respective modifications to undergo either strain-promoted azide-alkyne cycloaddition (SPAAC) or inverse-electron-demand Diels-Alder (IE-DA) reactions. We furthermore show that SPAAC is compatible with a previously published peptide-based attachment approach. By expanding the available toolkit to tag-free methods based on bioorthogonal reactions, we hope to enable researchers to interrogate the mechanics of a much broader range of proteins than is currently possible.

## Introduction

Mechanical forces are involved in varied biological processes such as force-bearing proteins in the muscle and tension upon chromosomes separation during cell division, and disruption of the cell’s ability to sense the mechanical properties of its surroundings represents a hallmark of many diseases. It is clear, therefore, that specific proteins must be able to sense mechanical signals and convert them into biological responses, but determining how they do so presents some major challenges. One such challenge is that measurements on single molecules are required, as force cannot be applied to a bulk solution of protein molecules but rather to individual protein molecules, the response being highly dependent on how and where on the protein the force is applied. To make single-molecule force measurements using optical tweezers, the protein needs to be attached to DNA handles that are then bound between two polystyrene or silica beads (Figure 1). Functionalisation of the beads and introduction of compatible DNA modifications is relatively straightforward, e.g. using streptavidin-biotin or digoxigenin-antidigoxigenin interactions. The current bottle-neck in single-molecule force-spectroscopy (SMFS) of proteins is the site-specific attachment of DNA in the case of optical tweezers, or protein/peptide handles in the case of atomic force microscopy. Various methods for the bio-conjugation of specific functionalities in proteins have been established^1^, but only a subset are stable enough to find application in SMFS. The current choices are: (i) endogenous cysteines for thiol-/maleimide-based attachments^2^, (ii) fusions of protein tags (HaloTag, SNAP-tag, SpyTag/Catcher)^3^, or (iii) N- and/or C-terminal introduction of small peptide tags^3^. The introduction of large protein tags can pose a problem for the expression of the protein of interest. Small peptide tags, such as the ybbR-tag^4^ and the sortase-tag^5^ can be beneficial in these cases as they enable linking of protein to handles post-translationally. Using the 4’-phosphopantetheinyl transferase (Sfp) enzyme, coenzyme A(CoA)-bearing moieties can be covalently cross-linked to a serine in the ybbR-tag^6, 7^. For example, the ybbR-tag has been used to link proteins to DNA oligos for both optical tweezers^8^ and AFM^4, 9^. Nevertheless, the use of such peptides is limited to the N- and C-termini, as they can interfere with protein folding and/or stability when introduced at internal sites^3^. Moreover, sortase-mediated attachment is limited to one terminus at a time due to the possible formation of circular protein products when termini of multiple molecules are conjugated^10^.

**Figure 1.**
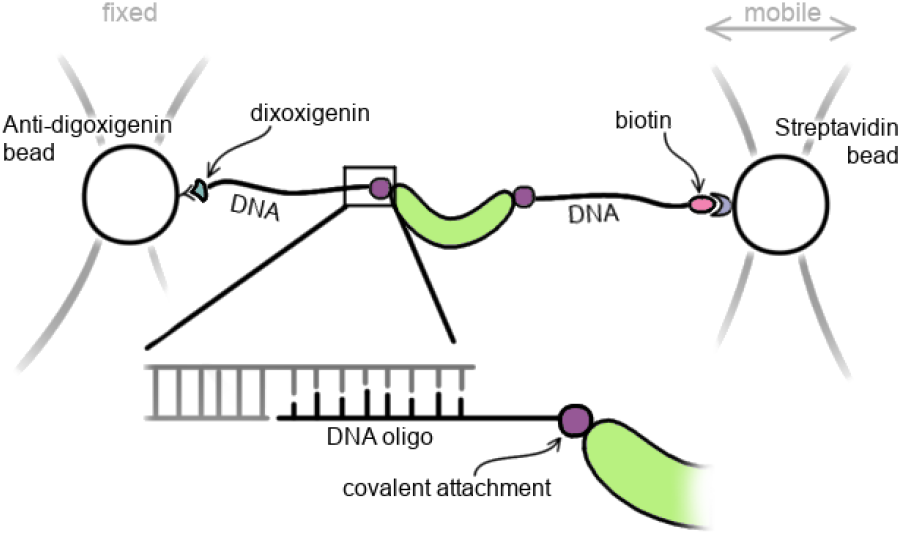
Site-specific attachment of DNA handles to a protein is necessary for force spectroscopy using a double-dumbbell optical tweezers set up. A small DNA oligo is covalently linked to the protein and hybridized to single-stranded overhangs in the DNA handles. The trapped beads can then be moved apart to exert a force on the protein.

Due to the necessity of site-specific modification, many of the most detailed force studies have used small ‘model’ proteins that lack amino acids (e.g. cysteine, lysine) with reactive side-chains that are most commonly used for conjugation reactions so that these amino acids can be engineered in at the desired sites. Current DNA-protein cross-linking protocol for optical tweezers experiments mainly use site-specific introduction of cysteine residues at either termini or internally of the protein of interest. After thiol-pyridine activation of cysteine side chains, proteins can then be conjugated to thiol-DNA oligomers^2^. However, unwanted by-products of this reaction can be poly-protein constructs and oligo homo-dimers, both of which are not easily separated from the protein-DNA chimera in purification steps following the conjugation reaction. Cysteine residues can also be reacted directly to maleimide-modified DNA oligos, but maleimide has to be supplied in large excess and the potential of maleimide dimerization remains^11, 12^. To improve reaction speeds and minimize the excess use of expensive components, Mukhortava and Schlierf^12^ developed a two-step protocol, in which cysteines are first functionalised with DBCO-maleimide followed by subsequent conjugation to azide-modified DNA oligos. Yet, these improvements are limited to proteins where cysteines are not present or where unwanted cysteines can be removed without severely affecting protein stability. For biology-driven research, however, the protein of interest rarely conforms to such requirements, and therefore there is tremendous need for new and broadly applicable approaches to site-specific conjugation.

Research in our group has focused on the folding and function of the giant HEAT-repeat protein PR65, the scaffold protein of the protein phosphatase 2A^13–15^. Up until now we were not able to conduct single molecule experiments to gain further insights due to the presence of a large number of cysteine residues in PR65 (14 in total) which complicate the specific attachment of dyes or DNA oligos. Previous experiments in our lab have shown that only a few of these can be substituted before protein stability is seriously affected. We therefore turned to a cysteine-independent method using different bio-orthogonal chemistries. These methods require the site-specific introduction of unnatural amino acids (UAAs) bearing desired chemical functionalities to react selectively with the corresponding modified DNA oligo. In addition, we designed protein constructs containing ybbR-tags and show that both attachment methods provide equivalent results in this case, and can be easily combined.

## Results

A variety of bioorthogonal chemistries have been developed for post-translational modification of bio-molecules which rely on the introduction of unnatural amino acids (UAAs) [reviewed extensively in 16, 17]. For SMFS experiments it is necessary that the functional chemical groups are inserted site-specifically. Therefore, we employed the amber codon suppression system developed by Chin and co-workers, which uses the amber codon tRNA (tRNA_*CUA*_) and aminoacyl tRNA synthetase (aaRS) from *Methanosarcina barkeri* combined with an orthogonal ribosome to incorporate pyrrolysine derivatives. In this manner, azide, alkyne and cyclopropene functionalities were introduced into PR65. These can then be reacted to DNA oligos bearing the necessary functionalities for either traditional “click” chemistry or copper-independent reactions (Figure 2).

**Figure 2.**
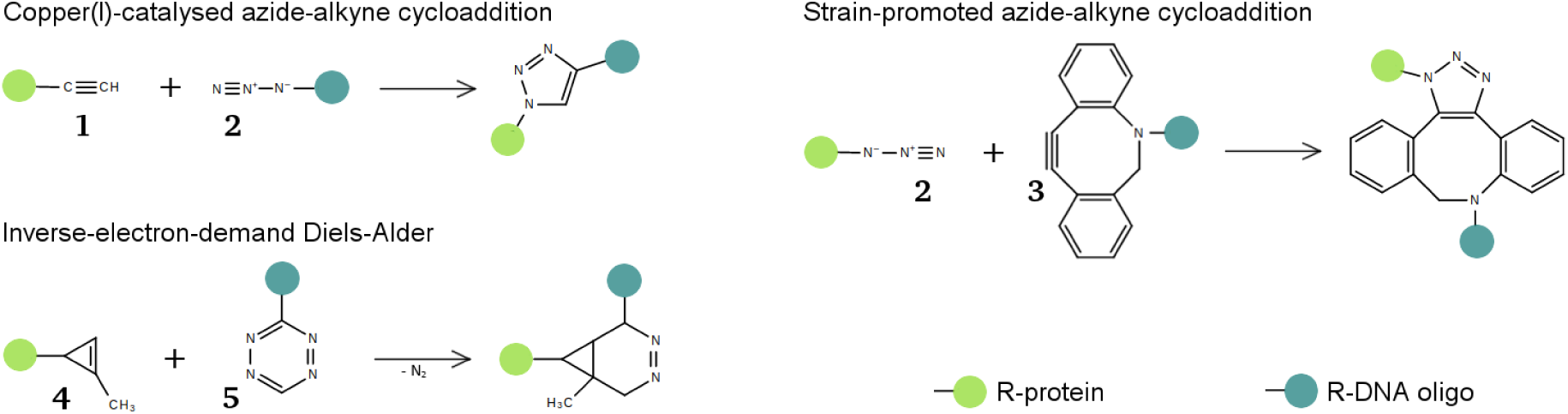
Bio-orthogonal reactions. Terminal alkynes (**1**) and azides (**2**) can form 1,4-disubstituted 1,2,3-triazoles in a 1,2-dipolar cycloaddition catalysed by copper. Azides can also react with strained alkynes such as DIBO/DBCO (**3**) in strain-promoted azide-alkyne cycloadditions that do not require a catalyst. Cyclopropenes (**4**) react with tetrazines (**5**) in inverse-electron-demand Diels-Alder reactions to diazanorcaradienes.

Amino acids in PR65 were replaced with UAAs as described previously^18–20^. Briefly, amber stop codons were introduced using site-directed mutagenesis before transferring the construct, bearing an N-terminal GST-tag and a C-terminal hexahistidine tag, into an expression vector containing an orthogonal 16S ribosomal subunit and the corresponding orthogonal ribosome binding sequence^18^. Due to its size and sequence repetition within the backbone, this vector poses some difficulties to the cloning process. After trialling different methods, we found that the In-Fusion cloning was the most reliable. The resulting construct was co-transformed into *E. coli* MDS42 ΔrecA together with a plasmid containing the pyrrolysine tRNA_*CUA*_ and aaRS^21^ and protein was expressed using 1 mM IPTG and 2 mM of UAA. PR65 was first purified by glutathione pull-down, and full-length product was separated from prematurely truncated protein by immobilized metal affinity chromatography. The protein yields per litre of culture were as expected for amber suppression of one or two codons, ranging from 0.5 mg/ml to 1.5 mg/l. We did not observe any correlation between the final yield and the type, number or location of the UAA. The incorporation of UAAs was verified by mass spectrometry of trypsin digests (Figure S1, Supplementary Information).

DNA oligos carrying a 3’-end azide modification were purchased. Oligos with DBCO and tetrazine modifications were produced as described previously using coupling reactions between amine-modified oligos and dual-functional small molecules, of which one group was an NHS-ester^9^. Free DBCO and tetrazine were removed by anion-exchange chromatography and modifications were verified by mass spectrometry (Figure S2, Supplementary Information). The modification of oligos with DBCO was successful, and no un-modified oligo could be detected. However, the modification of oligo with tetrazine did not proceed to completion, possibly due to hydrolysis of the NHS-ester. Attempts to separate the unmodified oligo from tetrazine-oligo by HPLC were unsuccessful, and hence subsequent reactions had to be carried out with this mixture. For screening purposes, reactions between protein and modified DNA oligos were generally carried out with the oligo present in equimolar quantity of the UAA, e.g. 5 µM protein bearing 2 UAAs were reacted with 10 µM modified oligo. The amount of final product can be increased by using DNA oligo in excess of the UAA. SDS-PAGE was used to analyse the reaction products as the added mass of the oligo (10 kDa) causes a noticeable retardation of the protein.

For end-to-end attachments, we exchanged residues D5 and L588 to amber stop codons. We first attempted copper-catalysed azide-alkyne cycloaddition (CuAAC) by incorporating an alkyne derivative of pyrrolysine into PR65, since azides incorporated during expression in *E. coli* could be altered in side reactions with endogenous thiols or by reducing agents^22^. Although the incorporation of alkyne pyrrolysing could be verified by mass spectrometry (Figure S1, Supplementary Information), we could only ever detect a small amount of protein conjugated to only one oligo, even when the azide oligo was present at 10-fold excess of the UAA and the reaction was incubated overnight at 37°C (Figure S3, Supplementary Information). In contrast, control reaction with 5-FAM-azide, a fluorescein derivative, were successful indicating that the alkyne moieties in PR65 were available for CuAAC. Further investigations using the azide oligo and 5-FAM-alkyne (Figure S4, Supplementary Information) led us to conclude that the oligo itself inhibits CuAAC in some manner, possibly by chelating Cu^2+^ and hence depleting the catalyst that is absolutely necessary for this reaction.

Copper is not required in strain-promoted azide-alkyne cycloadditions (SPAAC) and inverse-electron-demand Diels-Alder (IE-DA) reactions that require the introduction of an azide or cyclopropene into the protein, respectively. Purifications of azide-containing PR65 have to be carried out without reducing agents, but due to the relatively low yields after amber suppression, we could not detect a significant loss of protein due to intermolecular disulfide formation and subsequent aggregation. However, some reduction of azides to amines could be detected by mass spectrometry (Figure S5, Supplementary Information), whereas cyclopropene moieties remained intact. In SPAAC, products containing protein conjugated to one and two oligos are already detectable by SDS-PAGE after one hour (Figure 3a and Figure S6, Supplementary Information). The respective populations of un-modified, protein attached to one oligo and protein attached to two oligos shift with longer incubation times, reaching approximately 20%, 47% and 33%, respectively, after overnight incubation. Due to the presence of unmodified DNA oligo in the reaction mixture, IE-DA reactions were much slower and protein attached to two oligos can only barely be observed with coomassie stain after overnight incubation (Figure 3a).

**Figure 3.**
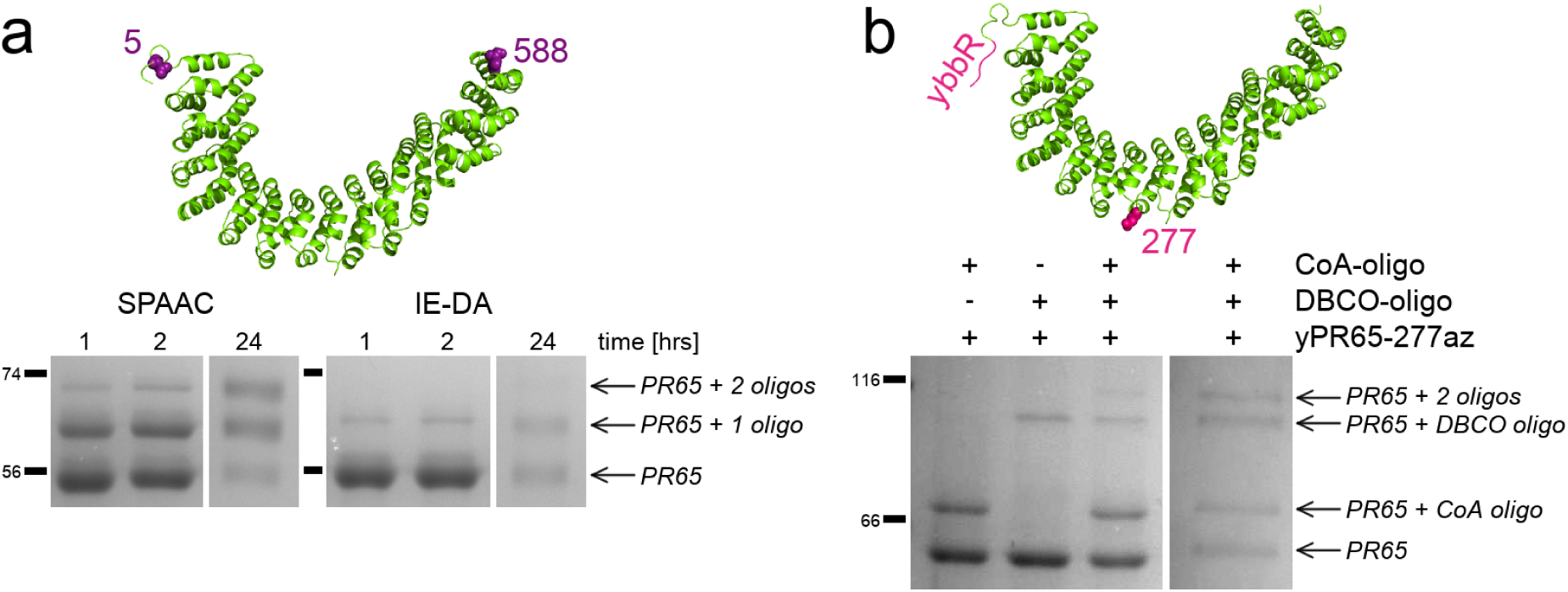
Attachment sites and SDS-PAGE of protein-oligo conjugations. (a) End-to-end attachment is achieved by substituting D5 and L588 with unnatural amino acids. When azides or cyclopropenes are incorporated and reacted with DBCO- or tetrazine-modified DNA oligos, reaction products can be visualised as larger molecular weight bands on an Coomassie stained SDS-PAGE gel. (b) Sfp-synthase mediated attachment and SPAAC are combined to exert forces on only the N-terminal half of the protein. Here, the first gel shows the reaction after 2 hours of incubation while the second shows the reaction containing both DNA oligos after overnight incubation. Un-cropped gels are available in the Supplementary Information.

In parallel with the bioorthogonal chemistry approaches, we also trialled the ybbR-tag system by directly fusing the sequence to the N- and C-terminus of PR65. When using the Sfp-enzyme at equimolar quantity of the CoA-modified DNA oligo, reaction efficiencies after overnight incubation were similar to SPAAC (Figure S7): approximately 38% of protein was attached to two oligos, 25% to one oligo and 36% remained unreacted. However, the ybbR-tag is a helical peptide^6^ and as such it could potentially alter the folding behaviour of the protein it is attached to. We performed chemical-induced equilibrium denaturation measurements of tagged and untagged PR65, and could not detect a significant difference beyond the usual experimental variation (Figure 4 and Table S1, Supplementary Information). By contrast, in a different protein context (consensus-designed tetra-tricopeptide repeats), we found that the ybbR-tag stabilized a 5-repeat array by 3.7 ± 0.6 kcal mol^−1^ and a 10-repeat array by 4.5 ± 0.9 kcal mol^−1^ (Figure S8). Moreover, the application of the ybbR-tag has limitations for the introduction of internal attachments sites, particularly in a repeat protein context where we have found that simple unstructured loops destabilize the array^23^. We expect that the introduction of a short α-helix would have an even greater effect, even if it is accompanied by unstructured spacers either side. Therefore, we used amber suppression for internal attachments, but combined it with an N-terminal ybbR-tag to demonstrate that Sfp-synthase mediated attachment of DNA is compatible with bioorthogonal chemical reactions. We introduced an N-terminal ybbR-tag and an internal amber codon at residue 277, which is located between the 7th and 8th HEAT repeats. The attachment of a DNA oligo at the internal site caused a retardation of around 30 kDa when analysed by SDS-PAGE, whereas migration of the N-terminal CoA-oligo attachment showed the expected retardation of 10 kDa (Figure 3 and Figure S9, Supplementary Information).

**Figure 4.**
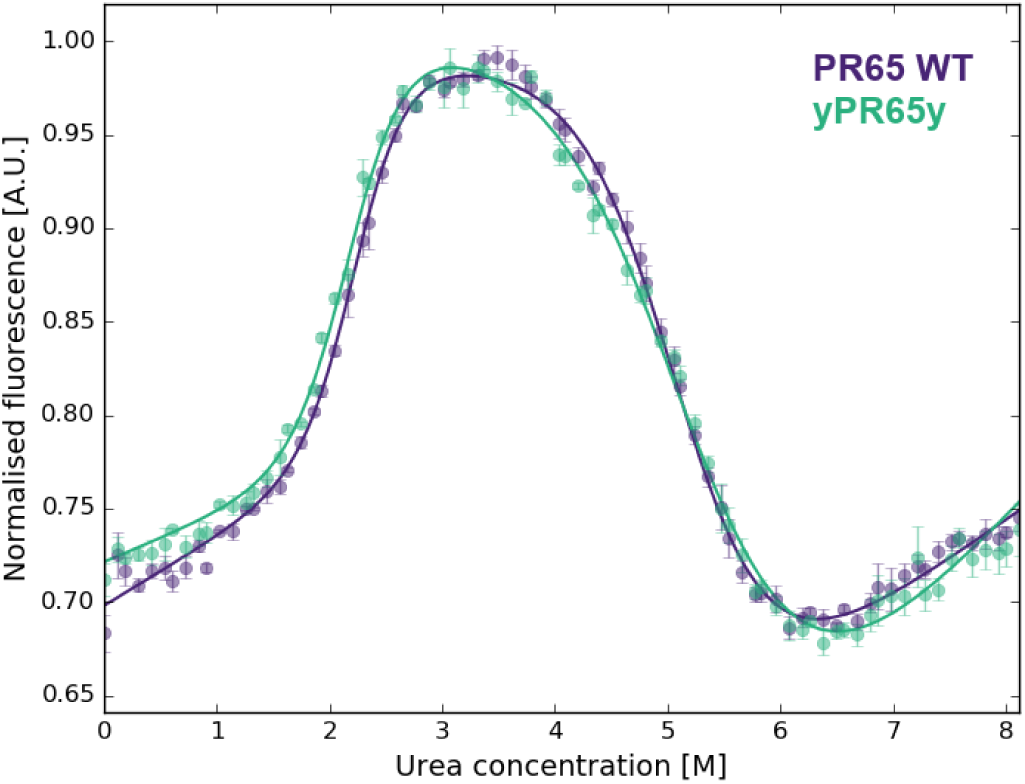
Equilibrium denaturations of PR65 WT (purple) and terminally ybbR-tagged PR65 (green). The data represent the average and standard error of the mean of three technical replicates for each experiment that were fitted with a three-state equation as described previously^15^. Parameters of the fit are listed in Table S1 in the Supplementary Information.

Since the reactions will contain a mixture of protein-DNA chimeras, further purification was performed using analytical grade FPLC (Figure S6, Supplementary Information) or HPLC (Figure S7, Supplementary Information) to remove unreacted protein and protein attached to one oligo only. Fractions containing protein attached to two oligos were mixed with a fixed amount of double-stranded DNA handles, that bear biotin and digoxigenin modifications at one end and a single stranded 5’-overhang at the other (Figure 1). The single-stranded overhang of the handles can then hybridise to the DNA oligos attached to the protein and the resulting hybridisation products can be analysed by agarose gel electrophoresis. The migration of DNA handles, which are approximately 550 bp in length, are retained when they are bound to protein (Figure 5). The hybridization of two DNA handles is dependent of the amount of protein present: the more protein is added the more the population distribution shifts towards protein bound to one handle only. At optimal protein concentration, mostly protein bound to two handles and unhybridised handles are present, the latter of which are filtered out manually during force spectroscopy experiments.

**Figure 5.**
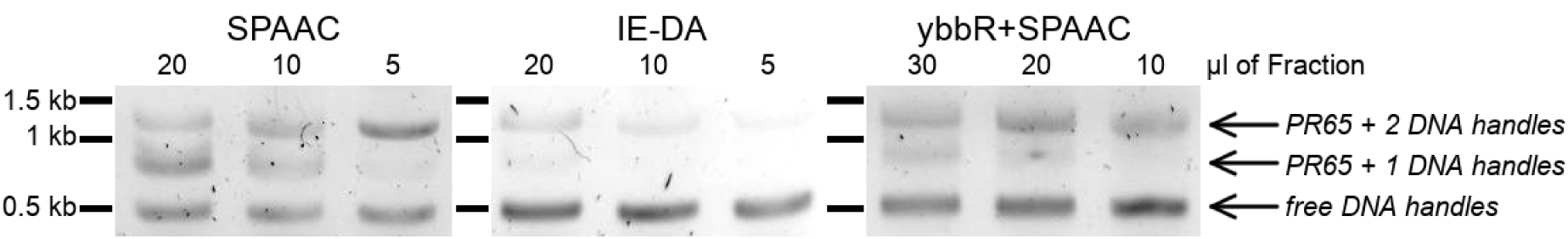
DNA handle hybridization to SPAAC, IE-DA and Sfp-mediated protein-oligo conjugations purified using an S200 10/300 GL column and detected by agarose gel electrophoresis. A protein with two DNA oligos attached can hybridize with one or two double stranded DNA handles. The amount of protein added to achieve maximal hybridisation with two handles is optimized by titrating protein while the DNA amount is kept constant. Un-cropped gels are available in the Supplementary Information (Figure S10).

Representative force-extension traces for the reported end-to-end attachments show PR65 to unfold in a series of three or four force peaks (Figure 6). The fully unfolded proteins are expected to have contour lengths of 200.41 nm and 202.21 nm for amber suppression and ybbR-tagged constructs, respectively. These theoretical values are calculated assuming 0.36 nm per amino acid, followed by subtraction of the physical dimensions of the protein structure as taken from crystal data (PDB ID 1b3u). We measured proteins of SPAAC attachments to have a contour length of 202 ± 3 nm (10 molecules), those of IE-DA attachments 194 ± 4 nm (8 molecules), and ybbR-based attachments 200 ± 5 nm (26 molecules).

**Figure 6.**
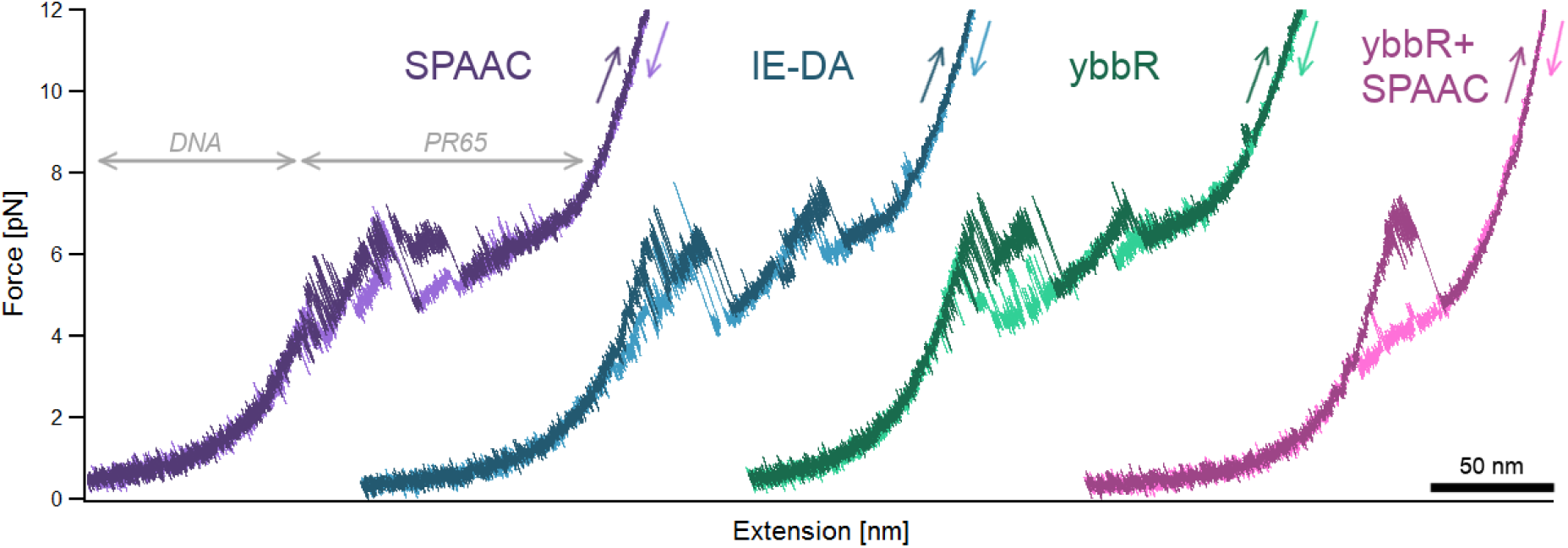
Representative full-length force-extension traces of all attachments. As a force is applied to the DNA-protein construct, the more pliable DNA stretches first before the protein experiences any force and unfolds in a series of force peaks combined with transitions that cannot be described with current polymer models. Unfolding and refolding traces are coloured in darker and lighter shades, respectively. All traces were obtained at pulling speeds of 10 nm s^−1^. See Figure S11 and S12 for multiple pulling traces of the same molecule.

The internal attachment, in which less than half of PR65 is unfolded, was measured to have a contour length of 84 ± 2 nm (10 molecules), although we expected a contour length of 93.32. This discrepancy is likely due to a repeat or some helices close to the C-terminal half of the protein remaining folded. The detailed discussion of the unfolding and refolding behaviour of PR65 is beyond the scope of this paper and will be published elsewhere. The variation observed between the three force curves of end-to-end attachment is characteristic of the protein system and independent of the attachment method (for further examples see Figures S11 and S12, Supplementary Information).

## Discussion

The amber suppression system was implemented successfully to produce PR65 with pyrrolysine derivatives carrying alkyne, azide and cyclopropene functional groups. Even though the purification of azide-containing PR65 was performed without DTT, a significant reduction from azide to amine could be detected using mass spectrometry, highlighting how important it is to further the development of site-specific bioorthogonal reactions involving functional groups that remain unaffected by reducing agents. In the future, the use of alkyne-bearing pyrrolysine will be limited to dye labelling reactions, which are as fast as previously reported^19, 20^, due to the interference of the oligo with CuAAC. Currently, the use of cyclopropene modifications for force-spectroscopy is limited by the production of sufficient tetrazine-functionalised DNA oligo. Mixtures of unconjugated and tetrazine-modified oligo were reacted to cyclopropene-containing PR65 with some success, albeit it decreased the yield of the final product significantly. However, this may be overcome by modifying the DNA-oligo with tetrazines using a different chemistry than that based on reacting amines with an NHS-ester. The Sfp-synthase-mediated conjugation of CoA to a ybbR-tag was explored as an alternative to bio-orthogonal chemistries. Both SPAAC and Sfp-reactions are similarly fast as long as Sfp-synthase is used in µM concentrations, and these two attachment methods can be combined with ease.

Depending on the application, both bioorthogonal chemistry and ybbR-tag based attachments have their advantages and disadvantages. Bioorthogonal chemistry allows site-specific modification of proteins without requiring the deletion of un-wanted cysteines or introduction of protein or peptide tags. Numerous chemistries are possible with different tRNA and aaRS systems, and only a few have been explored here. A disadvantage of using UAAs is that they can be expensive and the resulting protein yields are reduced significantly. However, given that single-molecule force spectroscopy requires relatively small quantities of protein, the yields obtained from just one litre of bacterial culture were more than sufficient for optimization and scale-up of the protein-DNA conjugation reaction. In contrast, Sfp-mediated site-specific modifications require the introduction of a small 11-residue tag which was not observed to interfere with protein expression. Sfp-synthase itself can be produced easily and cheaply in very high yields, and reducing agents do not interfere with the reaction. Just like any other tagging process, the effect of ybbR tagging on the stability of the protein will need to be examined. This check may be particularly important given the intrinsic helical propensity of the ybbR peptide. In conclusion, using both SPAAC and IED-DA, as well as Sfp-mediated reactions, we were able to generate protein-DNA conjugates. The methods described here therefore extend the toolbox available to scientists for the interrogation of single protein molecules by force.

## Methods

### Genetic constructs and mutagenesis

pRSF-oRibo-Q1-oGST-CaM_1TAG_^19^, containing an orthogonal ribosome under an IPTG-inducible promoter and the protein of interest under a constitutively active promoter, and pKW1^21^, containing the orthogonal aaRS and tRNA, for amber suppression were a kind gift from the Chin Lab (MRC LMB, Cambridge, UK). The PR65 template was available as a thrombin cleavable GST-PR65-H_6_ fusion protein in a pRSETa backbone. All primers are listed in Table S2 (Supplementary Information). The correct length and sequence of all constructs was verified by Sanger sequencing and restriction digests.

For amber suppression, constructs were created by first introducing the TAG codon at the positions of D5/L588 and E277 into the GST-PR65-H6 fusion protein using Round-the-Horn site-directed mutagenesis (RTH-SDM)^24, 25^. Different end-to-end attachment sites were trialled initially, but D5/L588 was the first clone to work and hence was carried forward. 100 µM primers were phosphorylated using polynucleotide kinase (ThermoFischer) and 2-3 mM ATP according to the manufacturers protocol. The enzyme was heat-inactivated at 85°C for 10-15min and phosphorylated primers were stored at −20°C until used. PCRs were performed using these primers and Phusion High-Fidelity DNA polymerase (NEB). PCR products were gel-purified and µg of DNA material was added to 1 µl Anza™T4 DNA Ligase Master Mix (ThermoFischer) in a total volume of 4 µl, incubated for 10-20 min at room temperature and transformed into in-house produced, chemically-competent DH5α *E. coli* cells.

The resulting constructs were then transferred into pRSF-oRibo-Q1-oGST by using the GST-internal SwaI and post-H_6_ SpeI restriction sites and In-Fusion Cloning (Takara Bio). pRSF-oRibo-Q1-oGST was digested using SwaI and SpeI, while the PR65 insert was obtained by PCR with primers that had 15bp overlap with these restriction sites and the vector backbone. Both vector and insert were gel purified and 1 µl of each was mixed with 0.5 µl 5X In-Fusion HD Enzyme Premix on ice. The reaction was incubated for 15 min at 50°Cin a pre-heated thermal cycler and placed back on ice immediately after. 2-4 µl of the ligation reaction were transformed into high efficiency DH5α *E. coli* (NEB).

N- and C-terminal ybbR-tags (DSLEFIASKLA) were introduced between thrombin cleavage site and M1, and A589 and the stop codon using RTH-SDM with primers bearing the tag sequence in the 5’ overhang.

### Protein expression and purification

PR65 WT and the ybbR-tagged version (yPR65y) were transformed into chemically competent C41 *E. coli* (Kommander laboratory, MRC-LMB, Cambridge). Suspension cultures were grown at 37°C in 2xYT media containing 50 µg/ml Ampicillin, shaking at 200 rpm until an OD_600_ of 0.6 to 0.8 was reached. Protein expression was induced with 250 µM isopropyl β-D-1-thiogalactopyranoside (IPTG, Generon) at 25°C overnight. Cells were harvested by centrifugation at 4000xg for 10 min at 4°C, before re-suspending in lysis buffer (50 mM Tris-HCl pH 7.5, 500 mM NaCl, 2 mM DTT) supplemented with EDTA-free protease inhibitor cocktail (Sigma Aldrich), and DNase I (Sigma Aldrich). The cells were lysed by passing the suspension through an Emulsiflex-C5 (AVESTIN) at pressures between 10000 and 15000 psi. Soluble protein was separated from cell debris and other insoluble fractions by centrifugation at 35000xg for 35 min at 4°C. The soluble protein fraction was applied to glutathione resin (Amintra Affinity Resins, Expedeon) equilibrated in lysis buffer. Resin was incubated at 4°C for 1-2 hours with rotation and cleaned using wash buffer (50 mM Tris-HCl pH 7.5, 150 mM NaCl, 2 mM DTT), followed by on-matrix cleavage of the protein using 50 units of bovine thrombin (Sigma) per litre of culture at 4°C overnight. Cleaved protein was removed from the resin using wash buffer. All fractions containing protein were pooled, diluted such that the NaCl concentration was < 50 mM and applied to a Mono Q 10/100 GL (GE Healthcare) equilibrated in 100 mM Tris-HCl pH 8.0, 2 mM EDTA, 0.5 g/l EGTA, 2 mM DTT. After washing the column, the protein was subsequently eluted using a 20 column volume salt gradient from 0 to 1 M NaCl. If necessary, MonoQ fractions containing the protein were concentrated before application to a HiLoad 26/600 Superdex 200 pg (GE Healthcare) equilibrated in PBS pH 7.4, 2 mM DTT. Fractions containing pure protein were pooled and concentrated using a Vivaspin®centrifugal concentrator

Expression and purification of amber suppression constructs was adapted from a protocol described by Sachdeva *et al.*^20^. GST-fusion proteins of PR65_5/588TAG_ and yPR65_277TAG_ were expressed in electro-competent MDS42 ΔrecA *E. coli* cells, grown at 37°C in 2xYT containing 25 µg/ml Kanamycin and 37.5 µg/ml Spectinomycin. Expression was induced at 37°C for 5 hrs when OD_600_ = 0.5-0.6, using 1 mM IPTG and 2 mM of either N-*ε*-(Prop-2-ynyloxycarbonyl)-L-lysine (Iris Biotech GmbH), N-*ε*-((2-Azidoethoxy)carbonyl)-L-lysine (Iris Biotech GmbH) or N-*ε*-[[(2-methyl-2-cyclopropene-1-yl) methoxy] carbonyl]-L-lysine (Sirius Fine Chemicals), which were dissolved in 0.2 M NaOH, diluted 1:3 using 1M HEPES pH 7.4 and adjusted to the pH of the cell culture. Proteins containing azides and cyclopropene derivatives were purified in buffers without reducing agent. All proteins were first purified by glutathione pull-down and thrombin cleavage at 4°C as described above. The cleavage product was applied to 1 ml of Ni-NTA resin per litre of culture (Amintra Affinity Resins, Expedeon) or a 1 ml HisTrap Excel column (GE Healthcare) equilibrated in wash buffer (50 mM Tris-HCl pH 7.5, 150 mM NaCl, optionally with 2 mM DTT). The resin was incubated at 4°C for 1-2 hours with rotation and cleaned with wash buffer, whereas column-bound protein was washed with 20 column volumes of wash buffer. Bound protein was recovered using elution buffer (100 mM Tris-HCl pH 8.0, 2 mM EDTA, 300 mM imidazole, 2 mM DTT). After analysis of the elution fractions obtained from either method by SDS-PAGE, the elutions were concentrated and buffer exchanged into PBS pH 7.4 with 1 mM DTT (alkyne) or without DTT (azide, cyclopropene) using Zeba Spin Desalting Columns (ThermoFisher Scientific).

For the expression of Sfp-synthase and CTPR proteins, please refer to the Supplementary Information. All proteins were flash-frozen in liquid N_2_ and stored at −80°C.

### Chemically modified DNA oligomers

DNA oligos for DNA-protein conjugations (Table S2) bearing 3’-end modifications with co-enzyme A or azide were acquired from Biomers and Integrated DNA Technologies, respectively. The modified oligos were resuspended in MilliQ H_2_O, aliquoted and stored at −20°C (azide, amine) or −80°C (CoA).

The protocol for chemical modification of oligos with DBCO and tetrazine functionalities was adapted from Nojima *et al.*^9^. DBCO-PEG_4_-NHS-ester (Sigma) and 6-methyl-tetrazine-PEG_5_-NHS-ester (Jena Bioscience, Figure S12, Supplementary Information) were conjugated to 3’-amine modified DNA oligos (Integrated DNA Technologies) in 50 µl Bicine-KOH pH 8.0 containing 100 µM amine and 5 mM NHS-ester. Due to its hydrophobicity, reactions containing DBCO were performed in 25% DMSO (Sigma) to ensure full solubility of the compound. The reaction was incubated at 37°C for 2-3 hours on an orbital shaker and loaded onto an anion-exchange colum (1 ml DEAE FF, GE Healthcare) equilibrated in 50 mM Tris-HCl pH 7.4. Bound oligo was eluted in one step using 50 mM Tris-HCl pH 7.4, 1M NaCl. The fractions containing oligo were isolated, split into 500 µl aliquots, combined with 1 ml ice cold, absolute ethanol and incubated at −80°C for at least 1 hour. The precipitate was pelleted for 30 min by centrifugation at 0°C and 20,000xg. The supernatant was carefully aspirated and discarded. The pellets were washed using 1 ml of room temperature, 95% (v/v) ethanol and collected by renewed centrifugation for 10 min at 4°C and 20,000xg. The supernatant was discarded, pellets were air dried and then resuspended in MilliQ H_2_O to be stored at −20°C.

### Conjugating DNA and protein

For optimization purposes CuAAC was performed in 20 µl volumes of PBS with 5 µM of alkyne bearing protein which were reacted to 100 µM azide using a range of catalyst concentrations. Copper sulfate (CuSO_4_), sodium ascorbate (NaAsc) and THPTA were pre-mixed into a “click mix” (CM). The 100X CM as defined by Sachdeva *et al.*^20^ contains 10 mM CuSO_4_, 25 mM NaAsc, 50 mM THPTA and was used in a final concentration of 1X. A 100X click mix with ten times the amount of NaAsc (CM-A) contains 10 mM CuSO_4_, 250 mM NaAsc and 50 mM THPTA^26^. Samples were incubated at 25°C for 0.5-2 hrs, or overnight. To stop the reaction the sample was either buffer exchanged or mixed directly with SDS-PAGE sample buffer. For proof of concept experiments and optimization, alkyne-bearing PR65 (alkPR65alk) was reacted with 5-FAM-azide (Lumiprobe). To produce protein-DNA chimeras, azide-functionalised DNA oligomers (Integrated DNA Technologies) were reacted with alkPR65alk. To test the functionality of the purchased azide-oligo, 20 µM of oligo was labelled with 20 µM 5-FAM alkyne dye (Lumiprobe) and increasing amounts of CM-A, sampling final CuSO_4_ concentration of 20 µM to 1 mM. These optimized conditions were then used to react 20 µM of protein with 100 µM azide oligo in a 20 µl volume using 10X CM-A.

SPAAC and IED-DA trial reactions were performed in 10 and 20 µl volumes of PBS containing 5 µM proteins and 10-20 µM modified oligo respectively. Control reactions for azide-bearing proteins were performed using TAMRA-DBCO (Jena Bioscience). Reaction mixtures were incubated for varying durations (0.5 hours to overnight) at room temperature in an orbital shaker.

For Sfp-synthase mediated DNA-protein conjugation, the reaction conditions as described by Yin *et al.*^7^ (100 µl of 50 mM HEPES pH 7.5, 10 mM MgCl_2_, 5 µM ybbR-tagged protein, 5 µM biotin-CoA and 0.1 µM Sfp-enzyme) did not yield detectable DNA-protein conjugation. Various conditions were screened and it was found that the Sfp concentration was the limiting factor. When including the enzyme to at least an equal stochiometric amount as that of the ybbR-tag, reactions yielded the desired product. When combining SPAAC and Sfp-mediated attachments, 5 µM of PR65 constructs containing one ybbR-tag and one azide functionality were reacted to 10 µM of each CoA-oligo and DBCO-oligo in 10 µl of 50 mM HEPES pH 7.5, 10 mM MgCl_2_. Control reactions were performed by omitting one oligo at a time.

Reaction products of oligo-dye couplings were analysed by electrophoresis using 1% unstained agarose gels. Protein-dye and protein-oligo reactions were analysed by SDS-PAGE. Before polyacrylamide gels were stained with Coomassie Blue, fluorescent bands were imaged under UV using a trans-illuminator (UVP, LLC).

Successful reactions were scaled up to 50 µl, containing 10 µM and a 1- or 2-fold excess (depending on availability) of the modified oligo with respect to the number of ybbR-tags and/or UAAs. After over-night incubation, these reactions were purified by size exclusion chromatography using either a Superdex 200 10/300 GL (GE Healthcare) or a YMC-Pack Diol-300 (Yamamura Chemical Research). Fractions were analysed by SDS-PAGE and those containing the majority of protein conjugated to two DNA oligos were hybridized to DNA handles and analysed by agarose gel electrophoresis.

### Equilibrium denaturations

WT and ybbR-tagged PR65 was buffer exchanged into 50 mM MES pH 6.5, 1 mM DTT. Samples of a total volume of 150 µl were prepared in black 96-well plates (Corning, low-binding) with urea gradients of 0 M to 8 M. The final protein concentrations were approximately 1 µM. Samples were incubated on an orbital shaker at 25°C for 2h. Tryptophans were excited at 295± nm and the fluorescence was monitored at 340±10 nm using a CLARIOStar microplate reader (BMG Labtech). The data from 4 reads were averaged, then normalised and fitted to a three-state equation:

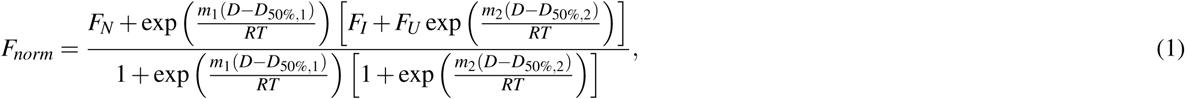

where *F*_*N*_ and *F*_*U*_ are the fluorescence of the folded and denatured states, respectively, and can be described by

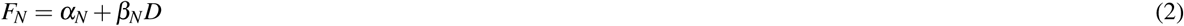

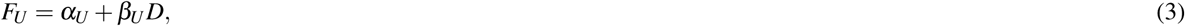

and *F*_*I*_, the fluorescence of the intermediates, is assumed to be constant. Subscripts of 1 and 2 refer to the first and second transitions, respectively.

### Force spectroscopy

Functionalised DNA handles of approximately 600 bp length were amplified from *λ* -DNA (e.g. Jena Bioscience) using the a triple biotinylated primer, a triple digoxigenin modified primer and a primer with a stable abasic site (Table S2, Metabion). A standard PCR was performed using *Taq* DNA polymerase in ThermoPol buffer (NEB) at an annealing temperature of 60°C and elongation temperature of 68°C. This PCR produces 5’-overhangs complementary to the oligo used for protein-DNA attachments. The final reaction was cleaned according to QIAquick PCR Purification Protocol with home made reagents and redissolved in water. The concentration was measured at 260 nm wavelength using a nano-photospectrometer (ThermoFischer).

Carboxyl-functionalised 1 µm silica beads (Bangs Laboratories) were modified in-house with anti-digoxigenin and tetramethylrhodamine-BSA (Sigma)^27^. Streptavidin coated beads were either produced in a similar manner or purchased (Bangs Laboratories). Anti-digoxigenin and streptavidin bead stocks were vortexed rigorously and diluted 1:20 and 1:130 in the appropriate buffer, respectively.

In general, 4-10 µl of purified DNA-protein construct were incubated with 100-200 ng of functionalised DNA handles for 0.5-1 hr at room temperature. Of that mixture, 0.5-3 µl were then incubated with 1 µl diluted anti-digoxigening beads in 10 µl sample buffer for no longer than 5 min. Finally, 0.5-0.7 µl of this mixture were added to 50 µl buffer containing 0.5-0.6 µl streptavidin beads, an oxygen scavenger system consisting of 0.65% (w/v) glucose (Sigma), 13 U ml^−1^ glucose oxidase (Sigma) and 8500 U ml^−1^ catalase (Calbiochem), and, if appropriate, 1-2 mM DTT or 1-5 mM TCEP.

Two parafilm strips were fixed to a microscope slide and covered by a cover slip. The combination is then heated to 80°C to melt the parafilm and thereby seal the chamber sides. The chamber is first blocked with 10 mg/ml BSA (Sigma) for 5 min and washed twice using the appropriate buffer before the sample is introduced. The edges are sealed with vacuum grease directly afterwards.

Experiments were conducted on a custom-built, dual-beam set up with back-focal plane detection^28^. All measurements presented in this work were performed in constant velocity mode, in which the mobile trap is moved away from the fixed trap at 10 nm s^−1^ to obtain force-extension traces. Trap stiffness ranged from 0.24 pN (commercial streptavidin beads) to 0.35 pN (in-house functionalised beads). Data were acquired at sampling rates of 20 kHz.

Force-extension data were analysed using the Igor software (WaveMetrics). Traces were fitted with worm-like chain (WLC) polymer models which describe the extension of DNA and a protein amino acid chain under force^29, 30^. The DNA force response can be described by the Modified Marko-Siggia WLC model^30^:

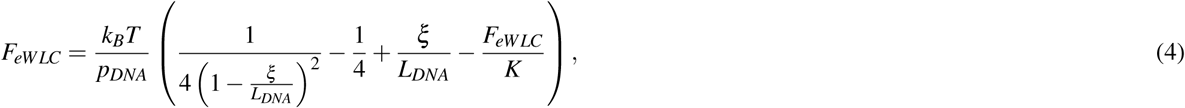

where *k*_*B*_ is the Boltzmann constant, *T* the temperature, *p*_*DNA*_ the persistence length of DNA, *L*_*DNA*_ the contour-length of the DNA and *K* its elastic stretch modulus. The protein force response can be modelled using the original Marko-Siggia WLC^29^:

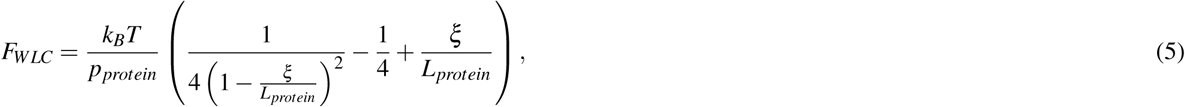

where *p*_*protein*_ = 0.7 is the persistence length of the protein and *L*_*protein*_ is the contour-length of the protein. The final extension of the protein-DNA construct is an addition of the stretching of both protein and DNA:

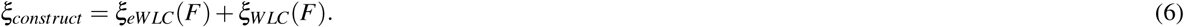

## Data availability

All data generated or analysed during this study are included in this published article and the corresponding Supplementary Information file. Raw data are available upon request. Plasmids for amber suppression can be obtained from the Chin laboratory (MRC-LMB, UK) after the completion of a Material Transfer Agreement. All other materials are available from the Itzhaki laboratory upon request.

## Acknowledgements

We thank Jason W. Chin and Kaihang Wang for helpful discussions on the introduction of unnatural amino acids. MS was funded by a studentship from the BBSRC Doctoral Training Partnerships, Cambridge. MR acknowledges funding through SFB 863 project A2 of Deutsche Forschungsgemeinschaft. LSI acknowledges the support of a Senior Fellowship from the UK Medical Research Foundation.

## Author contributions statement

MS and LSI designed the research. MS conducted the research, and DB and MR provided technical advice. All authors contributed to data analysis. MS and LSI wrote the manuscript text. All authors reviewed the manuscript.

## Additional information

The authors declare no competing interests.

## References

1. Stephanopoulos, N. & Francis, M. B. Choosing an effective protein bioconjugation strategy. Nat. Chem. Biol. 7, 876 (2011). Review Article.

2. Cecconi, C., Shank, E., Dahlquist, F., Marqusee, S. & Bustamante, C. Protein-DNA chimeras for single molecule mechanical folding studies with the optical tweezers. Eur. Biophys. J. 37, 729–738 (2008).

3. Ott, W., Jobst, M. A., Schoeler, C., Gaub, H. E. & Nash, M. A. Single-molecule force spectroscopy on polyproteins and receptor-ligand complexes: The current toolbox. J. Struct. Biol. 197, 3–12 (2017).

4. Pippig, D. A., Baumann, F., Strackharn, M., Aschenbrenner, D. & Gaub, H. E. Protein-DNA chimeras for nano assembly. ACS Nano 8, 6551–6555 (2014).

5. Durner, E., Ott, W., Nash, M. A. & Gaub, H. E. Post-Translational Sortase-Mediated Attachment of High-Strength Force Spectroscopy Handles. ACS Omega 2, 3064–3069 (2017).

6. Yin, J. et al. Genetically encoded short peptide tag for versatile protein labeling by Sfp phosphopantetheinyl transferase. Proc. Natl. Acad. Sci. 102, 15815–15820 (2005).

7. Yin, J., Lin, A. J., Golan, D. E. & Walsh, C. T. Site-specific protein labeling by Sfp phosphopantetheinyl transferase. Nat. Protoc. 1, 280 (2006).

8. Maillard, R. A. et al. ClpX(P) generates mechanical force to unfold and translocate its protein substrates. Cell 145, 459–469 (2011).

9. Nojima, T. et al. Nano-scale alignment of proteins on a flexible DNA backbone. PLoS ONE 7, 1–7 (2012).

10. Popp, M. W., Antos, J. M. & Ploegh, H. L. Site-specific protein labeling via sortase-mediated transpeptidation. Curr Protoc Protein Sci **Chapter 15**, Unit 15.3 (2009).

11. Jahn, M., Buchner, J., Hugel, T. & Rief, M. Folding and assembly of the large molecular machine Hsp90 studied in single-molecule experiments. Proc. Natl. Acad. Sci. 113, 1232–1237 (2016).

12. Mukhortava, A. & Schlierf, M. Efficient formation of site-specific protein-dna hybrids using copper-free click chemistry. Bioconjugate Chem. 27, 1559–1563 (2016).

13. Tsytlonok, M. & Itzhaki, L. S. Using FlAsH to probe conformational changes in a large HEAT repeat protein. Chem-BioChem 13, 1199–1205 (2012).

14. Tsytlonok, M., Sormanni, P., Rowling, P. J. E., Vendruscolo, M. & Itzhaki, L. S. Subdomain architecture and stability of a giant repeat protein. The J. Phys. Chem. B 117, 13029–13037 (2013).

15. Tsytlonok, M. et al. Complex energy landscape of a giant repeat protein. Structure 21, 1954–1965 (2013).

16. Lang, K. & Chin, J. W. Cellular incorporation of unnatural amino acids and bioorthogonal labeling of proteins. Chem. Rev. 114, 4764–4806 (2014).

17. Patterson, D. M., Nazarova, L. A. & Prescher, J. A. Finding the right (bioorthogonal) chemistry. ACS Chem. Biol. 9, 592–605 (2014).

18. Wang, K., Neumann, H., Peak-Chew, S. Y. & Chin, J. W. Evolved orthogonal ribosomes enhance the efficiency of synthetic genetic code expansion. Nat. biotechnology 25, 770–777 (2007).

19. Wang, K. et al. Optimized orthogonal translation of unnatural amino acids enables spontaneous protein double-labelling and FRET. Nat. Chem. 6, 393–403 (2014).

20. Sachdeva, A., Wang, K., Elliott, T. & Chin, J. W. Concerted, rapid, quantitative, and site-specific dual labeling of proteins. J. Am. Chem. Soc. 136, 7785–7788 (2014).

21. Rogerson, D. T. et al. Efficient genetic encoding of phosphoserine and its nonhydrolyzable analog. Nat. Chem. Biol. 11, 496–503 (2015).

22. Lang, K. & Chin, J. W. Bioorthogonal reactions for labeling proteins. ACS Chem. Biol. 9, 16–20 (2014).

23. Perez-Riba, A., Lowe, A. R., Main, E. R. G. & Itzhaki, L. S. Context-Dependent Energetics of Loop Extensions in a Family of Tandem-Repeat Proteins. Biophys. J. 114, 2552–2562 (2018).

24. Hemsley, A., Arnheim, N., Toney, M. D., Cortopassi, G. & Galas, D. J. A simple method for site-directed mutagenesis using the polymerase chain reaction. Nucleic Acids Res. 17, 6545–6551 (1989).

25. Moore, S. ’round the horn site-directed mutagenesis.

26. Hong, V., Presolski, S., Ma, C. & Finn, M. Analysis and optimization of copper-catalyzed azide-alkyne cycloaddition for bioconjugation. Angewandte Chemie Int. Ed. 48, 9879–9883 (2009).

27. Bauer, D. et al. A folding nucleus and minimal ATP binding domain of Hsp70 identified by single-molecule force spectroscopy. Proc. Natl. Acad. Sci. (2018).

28. von Hansen, Y., Mehlich, A., Pelz, B., Rief, M. & Netz, R. R. Auto- and cross-power spectral analysis of dual trap optical tweezer experiments using bayesian inference. Rev. Sci. Instruments 83, 095116 (2012).

29. Marko, J. F. & Siggia, E. D. Statistical mechanics of supercoiled DNA. Phys. Rev. E, Stat. Physics, Plasmas, Fluids Relat. Interdiscip. Top. 52, 2912–2938 (1995).

30. Wang, M. D., Yin, H., Landick, R., Gelles, J. & Block, S. M. Stretching DNA with optical tweezers. Biophys. J. 72, 1335–1346 (1997).

