## Supplementary Figures, Tables and Methods for "Bioorthogonal protein-DNA conjugation methods for force spectroscopy"

#### 1 Supporting figures

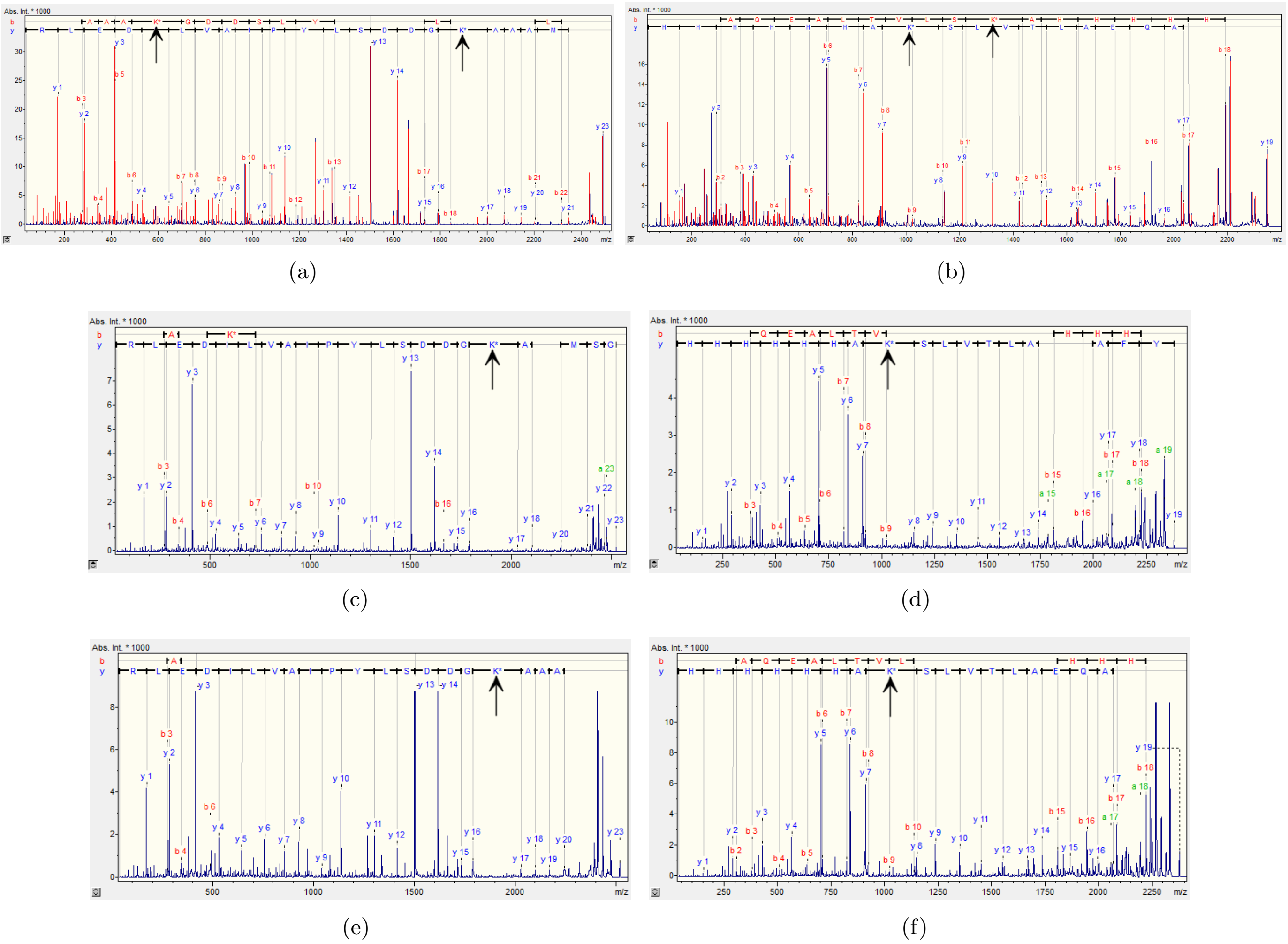


**Figure S1:** Tandem mass-spectrometry of PR65 peptides with UAAs incorporated at positions 5 and 588 (arrows). Shown are the N-/C-terminal peptides of alkPR65 (a,b), azPR65 (c,d), and cycPR65 (e,f). Data was provided by the PNAC Facility of the Biochemistry Department, University of Cambridge.

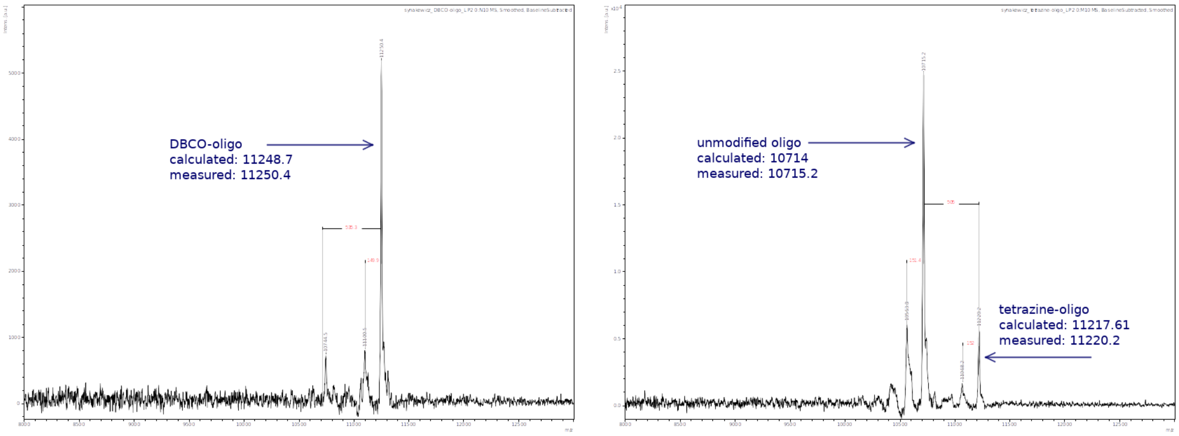


(a) (b)

**Figure S2:** MALDI mass-spectrometry of modified oligonucleotides (a) DBCO-oligo (b) tetrazine oligo. Measurements were performed by the PNAC Facility of the Biochemistry Department, University of Cambridge.

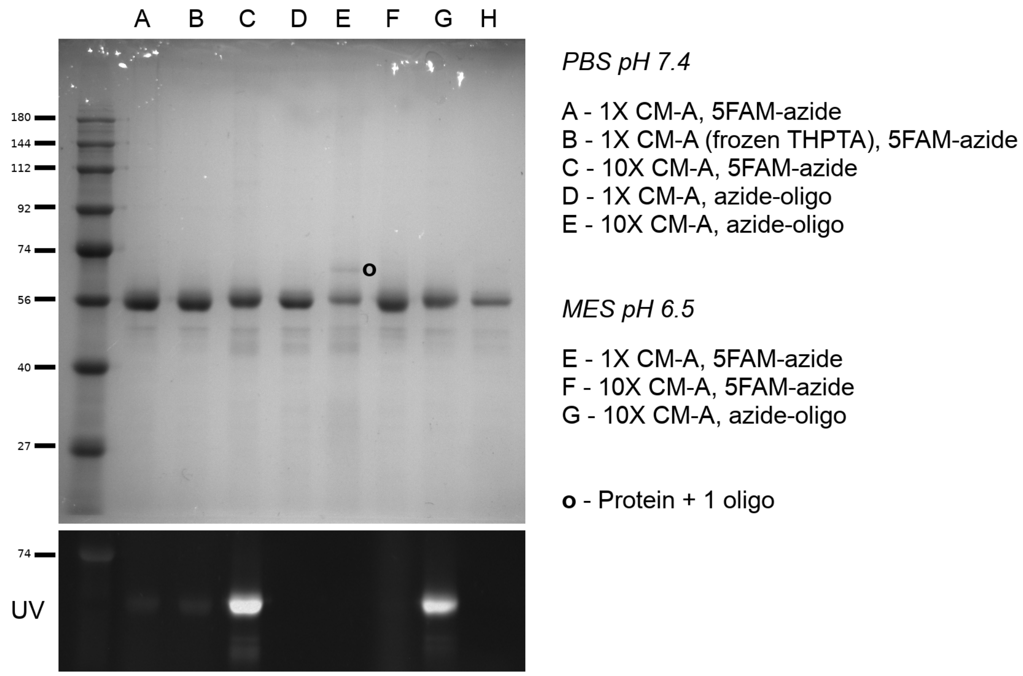


**Figure S3:** Example of click reactions using 100 µM 5-FAM azide or 3'-azide DNA oligo and 5 µ $PR65_{5/588}$ bearing alkyne moieties. Reactions were performed in PBS or MES buffer. Click-mix concentrations are given in fold excess of the standard CM-A concentration (click-mix containing 250 mM NaAsc).

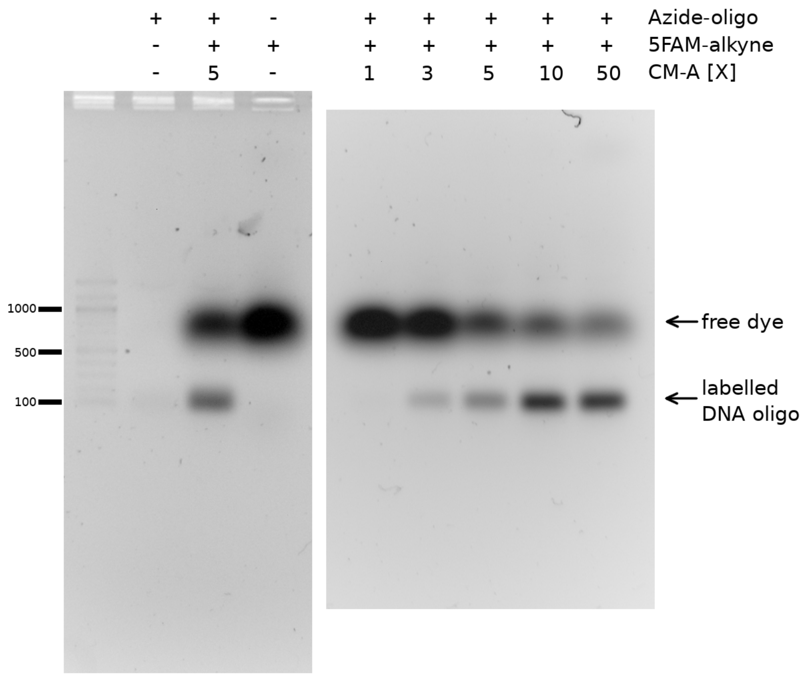


**Figure S4:** Click reactions of azide-oligo and 5-FAM-alkyne with increasing concentrations of CM-A (click-mix containing 250 mM NaAsc) on an unstained 1% agarose gel. Click-mix concentrations are given in fold excess of CM-A.

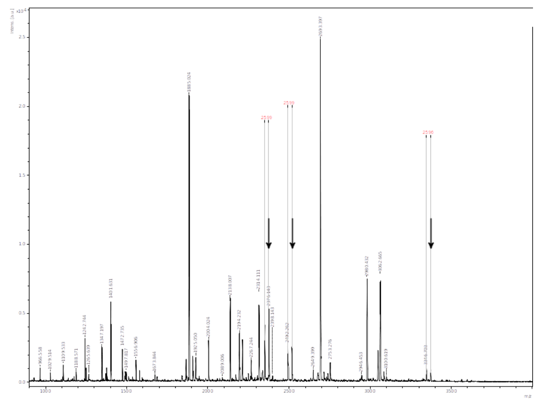

**Figure S5:** Each peptide containing azido-propargyllysine (arrows) is accompanied by a similar-size signal of -25.99amu resulting from the reduction of azide to amine. Measurements were performed by the PNAC Facility of the Biochemistry Department, University of Cambridge.

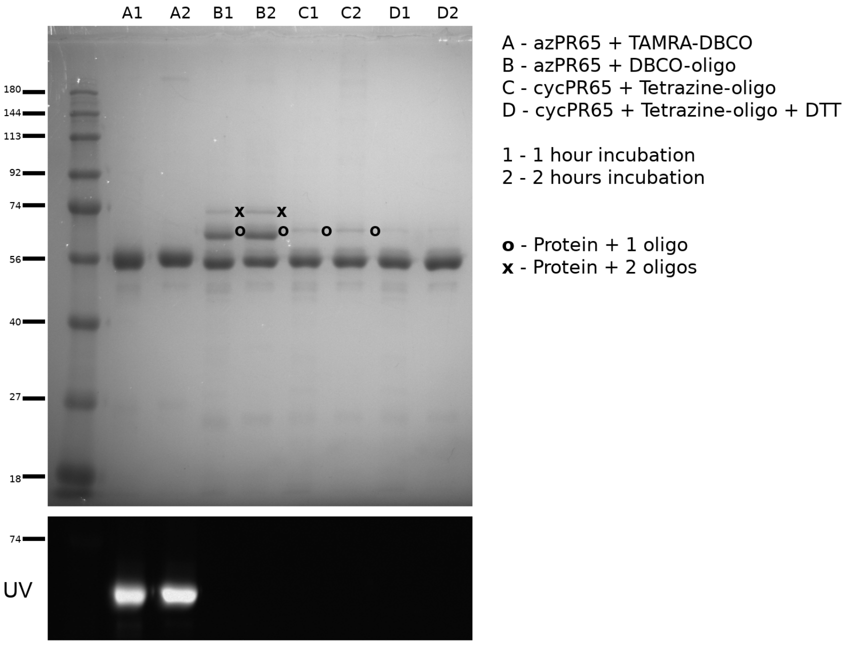


(a)

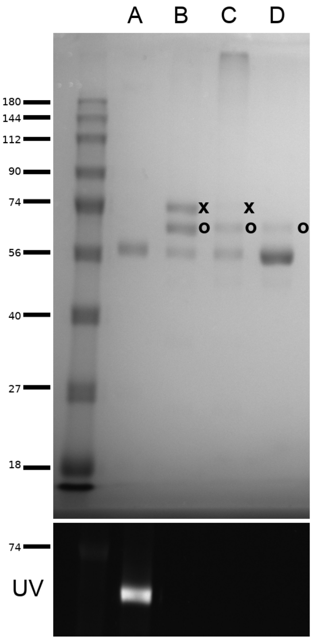


(b)

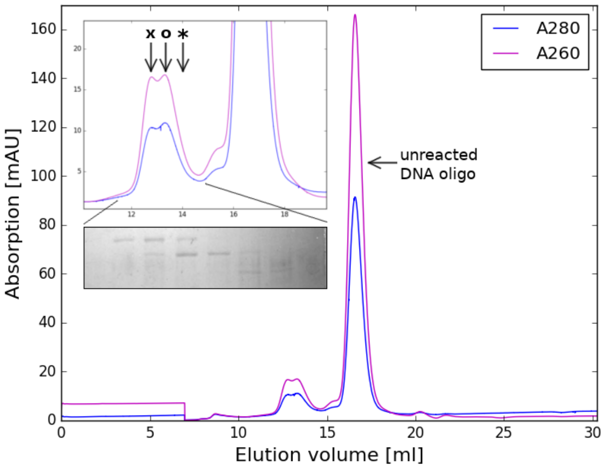


(c)

**Figure S6:** Full gels of Figure 3 in the main paper. SPAAC and IED-DA between azide or cyclopropene containing PR65$_{5/588\mathrm{TAG}}$ and TAMRA or oligo. (a) Example reactions using 5 µM protein and 20 µM TAMRA or oligo (b) Same reactions as in (a), incubated over night. (c) Chromatogram of a large scale reaction of 10 µM azPR65az and 40 µM DBCO-oligo incubated over night at 25°C, separated on a S200 10/300 GL. The smaller peak doublet preceding the oligo peak corresponds protein conjugated to two oligos (**x**) and one oligo (**o**). Un-reacted protein can be detected by SDS-PAGE even though a clear A280 peak is not visible (*****, inset).

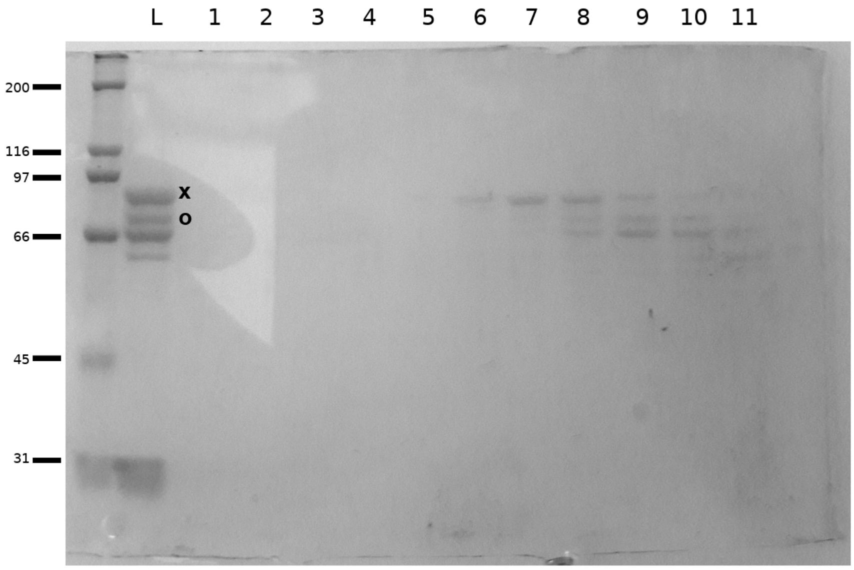


**Figure S7:** Sfp-synthase mediated attachment of 20 µM CoA-oligos to 10 µM of protein bearing N- and C-terminal ybbR-tags after an over-night incubation at room temperature (L). Lanes 1-11 are consecutive elution fractions after HPLC. Protein conjugated to two oligos (**x**) and one oligo (**o**) are present.

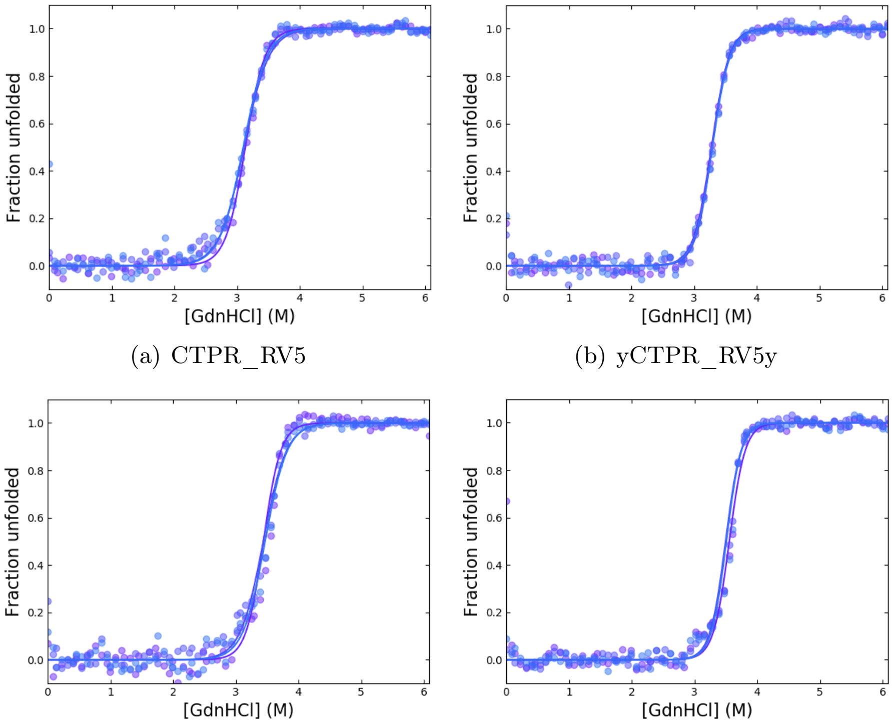


(a) CTPR_RV5 (b) yCTPR_RV5y

(c) CTPR_RV10 (d) yCTPR_RV10y

| Protein | $D_{50\%}$ [M] | $m$-value [kcal mol$^{-1}$ M$^{-1}$] | $\Delta G_{U-N}^{H_2O}$ [kcal mol$^{-1}$] |
|---|---|---|---|
| CTPR_RV5 | 3.114 ± 0.008 | 3.3 ± 0.2 | 10.3 ± 0.6 |
| yCTPR_RV5y | 3.287 ± 0.003 | 4.26 ± 0.07 | 14.0 ± 0.2 |
| CTPR_RV10 | 3.449 ± 0.007 | 3.6 ± 0.3 | 12.5 ± 0.9 |
| yCTPR_RV10y | 3.52 ± 0.01 | 4.82 ± 0.06 | 17.0 ± 0.2 |

**Figure S8:** Equilibrium denaturation using guanidinium hydrochloride of untagged CTPR_RV and ybbR-tagged CTPR_RV proteins with 5 and 10 repeats, respectively. The curves represent three technical replicates for each experiment that were fitted independently with a two-state equation. Due to the nature of the unfolding mechanism of CTPRs, which can be better described by Ising models [1], the parameters obtained from the two-state fit are only apparent values.

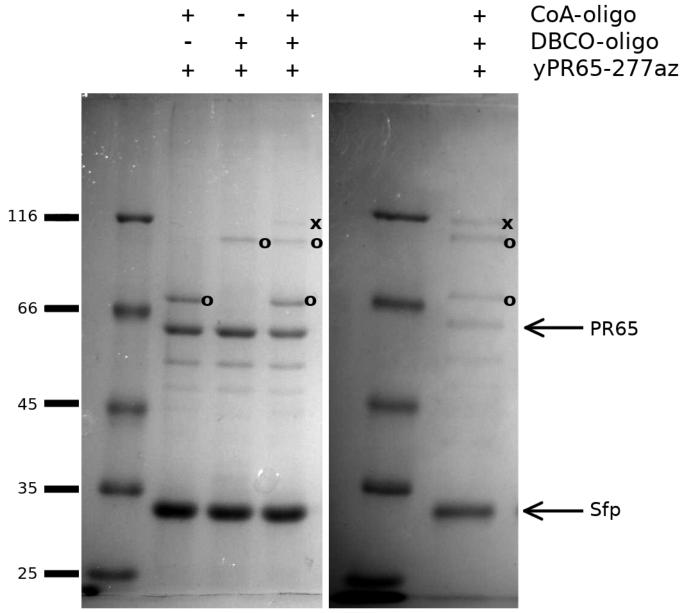


**Figure S9:** Combining CoA-ybbR and SPAAC conjugations. Conjugations of yPR65-277az to DBCO- and CoA oligonucleotides after 2 hours (left) and over-night incubation (right).

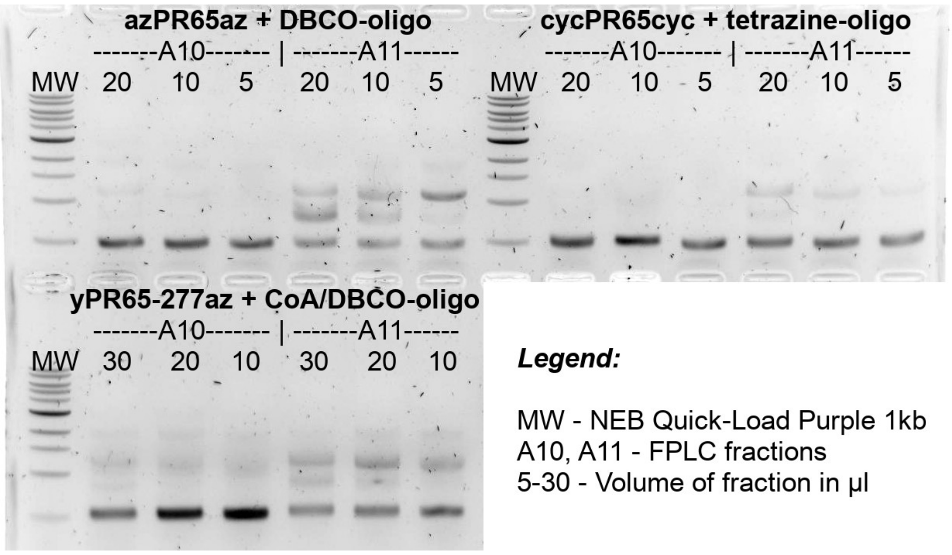


**Figure S10:** DNA handle hybridization to SPAAC, IE-DA and Sfp-mediated protein-oligo conjugations purified using an S200 10/300 GL column and detected by agarose gel electrophoresis. Shown are hybridization products between varying amounts of protein from two different fractions to double stranded DNA handles.

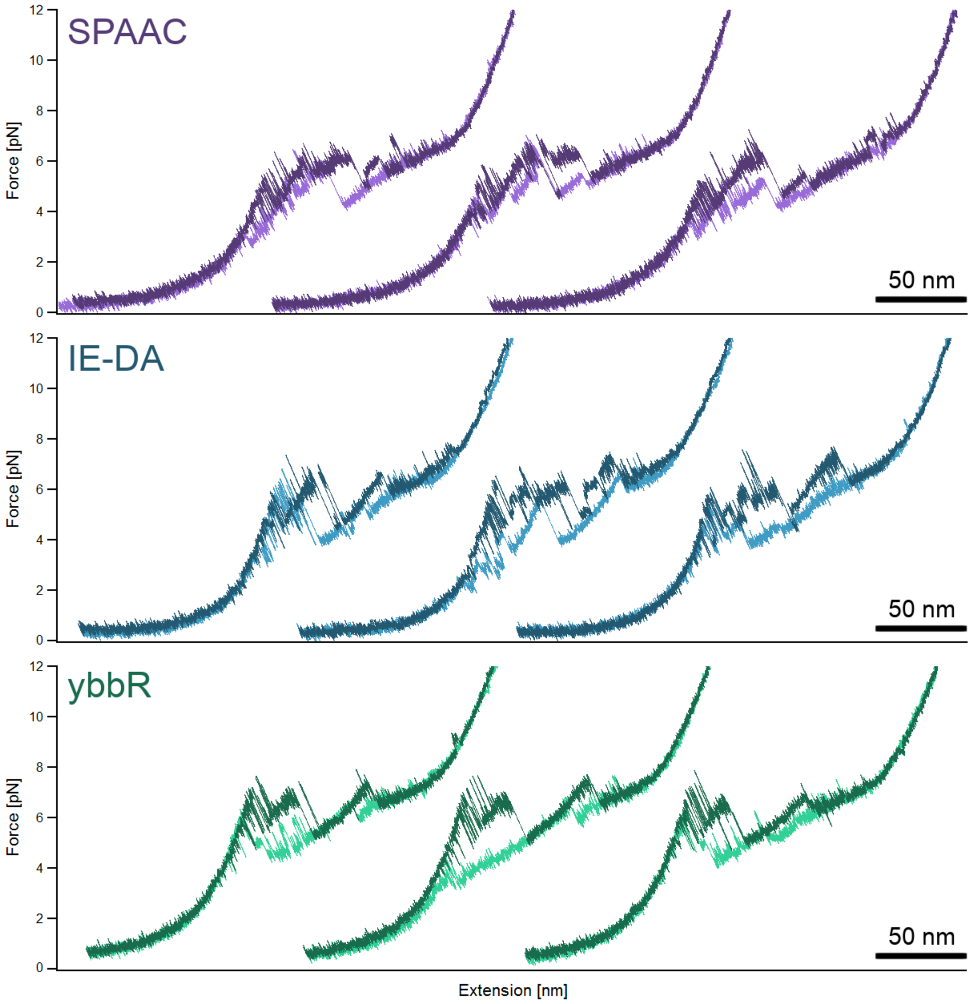


**Figure S11:** Representative full-length force extension curves of the same molecule for each PR65 attachments. Unfolding traces are coloured in darker shades, while refolding traces are coloured in lighter shades. All traces were taken at pulling speeds of 10 nm $s^{-1}$.

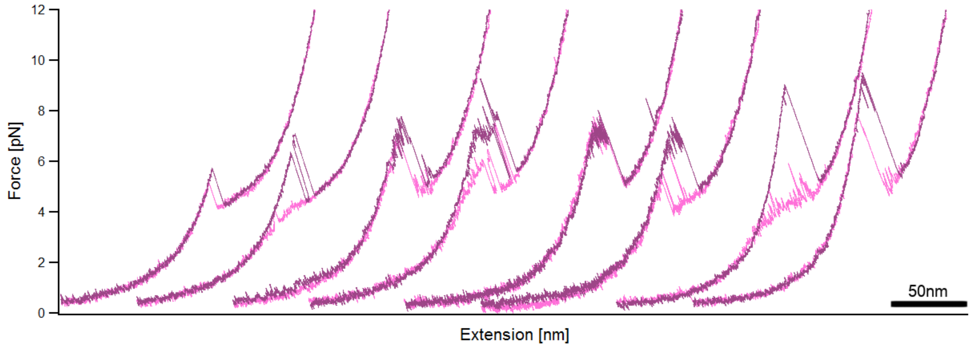


**Figure S12:** Two representative stretch-relax cycles of four yPR65-277az molecules. All data shown were obtained at pulling velocities of 10 nm $s^{-1}$.

### 2 Supporting tables

**Table S1:** Equilibrium denaturation three-state fit parameters of PR65 variants. The parameters represent averages ± s.e.m. of three technical replicates.

| Protein | $D_{50\%-1}$ [M] | $m_1$ [kcal mol$^{-1}$ M$^{-1}$] | $D_{50\%-2}$ [M] | $m_2$ [kcal mol$^{-1}$ M$^{-1}$] |
|---|---|---|---|---|
| PR65 | 2.24 ± 0.06 | 2.51 ± 0.09 | 5.13 ± 0.03 | 1.40 ± 0.02 |
| yPR65y | 2.14 ± 0.06 | 2.48 ± 0.06 | 5.20 ± 0.05 | 1.16 ± 0.06 |

**Table S2:** List of DNA oligonucleotides (5' to 3'). * indicates sites of modification

| Name | Sequence |
|---|---|
| PR65 D5TAG Fwd | TAGGGCGACGACTCGCTGTACCCC |
| PR65 D5TAG Rev | GGCCGCCGCCATATGATGATGATGG |
| PR65 L588TAG Fwd | TAGGCCCACCACCACCACCACC |
| PR65 L588TAG Rev | AGACAGAACAGTCAGAGCCTCCTGGGC |
| PR65 E277 Fwd | TAGATCACCAAGACAGACCTGGTC |
| PR65 E277 Rev | AGGCCCCACTGCTTTCTGG |
| oRibo Inf SwaI Fwd | TCATAAAACATATTTAAATGGTGATCATGTAACCCATCC |
| oRibo Inf SpeI Rev | GTACGGCCGACTAGTTCAGTGGTGGTG GTGGTGGTGGGCGAGAG |
| PR65 NybbR Fwd | GATTCTCTTGAATTTATTGCTAGTAAGC TTGCGATGGCGGCGGCCGACGGCGACG |
| PR65 NybbR Rev | GGATCCACGCGGAACCAGATCCGATTTTGG |
| PR65 CybbR Fwd | CACCACCACCACCACCACTGAAAGC |
| PR65 CybbR Rev | CGCAAGCTTACTAGCAATAAATTCAAGAG AATCGGCGAGAGACAGAACAGTCAGAGC |
| CTPR_RV NybbR Fwd | TGCTAGTAAACTTGCGGCAGAAGCAC TGAATAATCTGGG |
| CTPR_RV NybbR Rev | ATAAATTCAAGAGAATCGGATCCACGCGGAACCAG |
| CTPR_RV CybbR Fwd | TGCTAGTAAACTTGCGTAATAAAAGCTTGATCCGGC |
| CTPR_RV CybbR Rev | ATAAATTCAAGAGAATCAGATCTCGGGTCCAGTTCC |
| DNA-protein coupling | GGCAGGGCTGACGTTCAACCAGACCAGCGAGTCG* |
| Biotin/digoxigenin | *GGCGA*CTGG*CGTTGATTTG |
| Abasic | CGACTCGCTGGTCTGGTTGAACGTCAGCCC TGCC*CCTGCCCGGCTCTGGACAGG |

### 3 Supporting methods

#### 3.1 Genetic constructs and mutagenesis of CTPR constructs

DNA constructs of CTPR_RV proteins were build sequentially from from single/double repeat modules using BamHI/BglII cloning as previously described by Kajander *et al.* [1]. In brief, a single repeat in a pRSET backbone is preceded by a BamHI restriction site and followed by a BglII restriction site, double stop codon and HindIII restriction site. To create a construct containing $N$ repeats, a vector containing $M$ repeats is digested using BglII and HindIII and a vector containing a single repeat is amplified by PCR using Phusion DNA polymerase and T7-forward and -terminator primers. Both the vector restriction digest and the PCR product are gel purified. The PCR product is subsequently digested using BamHI and HindIII and the enzymes are heat-inactivated at 80°C for 10 min. Since BamHI and BglII produce the same 5'-overhangs the single repeat can be ligated into the vector using QuickStick ligase (Bioline). The ligation product should then contain $M+1$ repeats. The whole procedure is repeated until an $N$-repeat construct has been obtained.

To create ybbR-tagged CTPRs, the tag sequence was first introduced by RTH mutagenesis directly adjacent to the repeat sequence either N-terminally or C-terminally of a single repeat, giving rise to yCTPR_RV1 and CTPR_RV1y respectively. Second, the required number of repeats were added to yCTPR_RV1, resulting in yCTPR_RV4 and yCTPR_RV9. Last, the C-terminally tagged repeat was added to produce constructs with $N = 5, 10$ that contained both N- and C-terminal ybbR-tags.

All primers are listed in Table S2. The correct length and sequence of all constructs was verified by sanger sequencing and restriction digests.

#### 3.2 Protein expression and purification

##### 3.2.1 Sfp-synthase

The pCK plasmid containing Sfp-synthase-$H_6$ was a kind gift from the Gaub laboratory (Ludwig Maximilian Universität, München, Germany). For expression, the plasmid was transformed into chemically competent C41 *E. coli* and plated onto LB Agar containing Ampicillin. All colonies from the plate were used to inoculate 1 l of 2xYT. Protein expression was induced at $OD_{600} \sim 0.6$ by the addition of 1 mM IPTG and followed by further incubation at 25°C overnight. Following centrifugation, the pellet was resuspended in lysis buffer (20 mM Tris-HCl pH 7.5, 500 mM NaCl, 5 mM imidazole) containing protease inhibitor cocktail and DnaseI, lysed as described above and centrifuged again to remove cell debris. The soluble protein fraction was filtered through a 0.22 µm PES membrane and applied to a 5 ml HisTrap Excel column (GE Healthcare) equilibrated in lysis buffer. After washing with 20 column volumes of buffer, Sfp was eluted in one step using 20 mM Tris-HCl pH 8.0, 300 mM imidazole, 300 mM NaCl, 2 mM EDTA. The elution fractions were analysed by SDS-PAGE and those containing protein at >90% purity were pooled. This solution was then split in half and dialysed twice against either 10 mM Tris-HCl pH 7.5 or PBS pH 7.4 containing 1 mM EDTA and 10% (v/v) glycerol. Protein was flash-frozen after concentration using a Vivaspin®centrifugal concentrator and stored at -80°C.

##### 3.2.2 CTPRs

N-terminally $H_6$-tagged CTPR proteins were transformed in C41 *E. coli* and plated on LB Agar containing 100 µg $ml^{-1}$ Ampicillin. All colonies were used to inoculate 0.5 l of 2xYT. Protein expression was induced using 1 mM IPTG over 3 hours. After lysis in 50 mM Tris-HCl pH 7.5, 500 mM NaCl, 20 mM imidazole containing protease inhibitor cocktails and DnaseI, the cell suspension was heated to 70-80°C in a water bath before centrifugation. The soluble protein was then filtered through a 0.22 µm PES membrane and applied to a 5 ml HisTrap Excel column equilibrated in 50 mM Tris-HCl pH 7.5, 500 mM NaCl, 20 mM imidazole. The column was washed using 20 column volumes of buffer before proteins were eluted in 50 mM Tris-HCl pH 7.5, 150 mM NaCl, 300 mM imidazole. All fractions containing protein were pooled, and if necessary, concentrated to <15 ml total volume using a Vivaspin® centrifugal concentrator. The protein was then further purified by size exclusion chromatography using a HiLoad 26/600 Superdex 75 pg (GE Healthcare) equilibrated in 50 mM sodium phosphate pH 6.8, 150 mM NaCl. The fractions were analysed using SDS-PAGE and those containing the least amount of recombined protein were pooled and concentrated.

##### 3.3 Denaturing CTPR proteins using GdHCl

Samples of a total volume of 150 µl were prepared in a 96-well format (Greiner, medium-binding), in 50 mM sodium phosphate pH 6.8, 150 mM NaCl with guanidinium hydrochloride (GdHCl) gradients of 0 to 6 M. Final protein concentrations ranged from 0.3 µM for larger constructs to 10 µM for smaller constructs. Samples were incubated on an orbital shaker at 25°C for 2h. Tyrosine fluorescence was excited at $280 \pm 10$ nm and emission was monitored at $330 \pm 10$ nm using a CLARIOStar microplate reader (BMG Labtech). The data from 9 reads was averaged and normalised by the maximal fluorescence. Usually, these data would then be fitted to a two-state equation:

$$F_{norm} = \frac{\alpha_N + \beta_N D + (\alpha_U + \beta_U D)\exp(\frac{m(D-D_{50\%})}{RT})}{1 + \exp(\frac{m(D-D_{50\%})}{RT})}, \tag{1}$$

where $D$ is the denaturant concentration, $\alpha_N$ and $\alpha_U$ the fluorescence signal of the native and unfolded state at 0.0 M Urea, $\beta_N$ and $\beta_U$ are their respective rates of change with increasing denaturant, $D_{50\%}$ is the midpoint of the unfolding transition, and $m$ is the constant of proportionality related to the change in solvent accessible surface area upon unfolding. However, since the denaturation profile of larger CTPR arrays deviates from a simple-two state unfolding behaviour especially in the first half of the transition, such a fit can affect the apparent $m$-value by compensating for the deviation with the baseline parameters. Therefore, the baselines were fitted first using a straight-line equation to obtain values for $\alpha_N$, $\alpha_U$, $\beta_N$ and $\beta_U$. These were then fixed to obtain the apparent $D_{50\%}$ and $m$-value using Equation 1.

Assuming that all protein is folded at zero denaturant and fully unfolded at high denaturant concentrations, the fluorescence can be converted into fraction folded, $\theta$, or fraction unfolded, $1-\theta$, using

$$\theta = \frac{F - \alpha_U - \beta_U D}{\alpha_N - \alpha_U + (\beta_N - \beta_U)D}, \tag{2}$$

or

$$1 - \theta = 1 - \frac{F - \alpha_U - \beta_U D}{\alpha_N - \alpha_U + (\beta_N - \beta_U)D} = \frac{-F + \alpha_N + \beta_N D}{\alpha_N - \alpha_U + (\beta_N - \beta_U)D} \tag{3}$$

#### References


[1] T. Kajander, A. L. Cortajarena, E. R. G. Main, S. G. J. Mochrie, and L. Regan. A new folding paradigm for repeat proteins. *Journal of the American Chemical Society*, 127(29):10188–10190, 2005.

[2] K. Wang, A. Sachdeva, D. J. Cox, N. M. Wilf, K. Lang, S. Wallace, R. A. Mehl, and J. W. Chin. Optimized orthogonal translation of unnatural amino acids enables spontaneous protein double-labelling and FRET. *Nature Chemistry*, 6:393–403, 2014.

[3] D. T. Rogerson, A. Sachdeva, K. Wang, T. Haq, A. Kazlauskaite, S. M. Hancock, N. Huguenin-Dezot, M. M. Muqit, A. M. Fry, R. Bayliss, and J. W. Chin. Efficient genetic encoding of phosphoserine and its nonhydrolyzable analog. *Nature Chemical Biology*, 11(7):496–503, 2015.

[4] A. Hemsley, N. Arnheim, M. D. Toney, G. Cortopassi, and D. J. Galas. A simple method for site-directed mutagenesis using the polymerase chain reaction. *Nucleic Acids Research*, 17(16):6545–6551, 1989.

[5] S. Moore. 'round the horn site-directed mutagenesis. URL `https://openwetware.org/wiki/%27Round-the-horn_site-directed_mutagenesis`.

[6] A. Sachdeva, K. Wang, T. Elliott, and J. W. Chin. Concerted, rapid, quantitative, and site-specific dual labeling of proteins. *Journal of the American Chemical Society*, 136(22):7785–7788, 2014.

[7] T. Nojima, H. Konno, N. Kodera, K. Seio, H. Taguchi, and M. Yoshida. Nano-scale alignment of proteins on a flexible DNA backbone. *PLoS ONE*, 7(12):1–7, 2012.

[8] V. Hong, S. Presolski, C. Ma, and M. Finn. Analysis and optimization of copper-catalyzed azide-alkyne cycloaddition for bioconjugation. *Angewandte Chemie International Edition*, 48(52):9879–9883, 2009.

[9] J. Yin, A. J. Lin, D. E. Golan, and C. T. Walsh. Site-specific protein labeling by Sfp phosphopantetheinyl transferase. *Nature Protocols*, 1:280, 2006.

[10] D. Bauer, S. Meinhold, R. P. Jakob, J. Stigler, U. Merkel, T. Maier, M. Rief, and G. Žoldák. A folding nucleus and minimal ATP binding domain of Hsp70 identified by single-molecule force spectroscopy. *Proceedings of the National Academy of Sciences*, 2018. ISSN 0027-8424.

[11] Y. von Hansen, A. Mehlich, B. Pelz, M. Rief, and R. R. Netz. Auto- and cross-power spectral analysis of dual trap optical tweezer experiments using bayesian inference. *Review of Scientific Instruments*, 83 (9):095116, 2012.

[12] J. F. Marko and E. D. Siggia. Statistical mechanics of supercoiled DNA. *Physical Review E, Statistical Physics, Plasmas, Fluids and Related Interdisciplinary Topics*, 52(3):2912–2938, 1995.

[13] M. D. Wang, H. Yin, R. Landick, J. Gelles, and S. M. Block. Stretching DNA with optical tweezers. *Biophysical Journal*, 72(3):1335–1346, 1997.
